# *MUC4* regulatory-region methylation is remodeled during breast cancer progression and metastasis

**DOI:** 10.64898/2026.11.11.765423

**Authors:** Savannah R. Free, Liliana Loza Sanchez, Macey M. McAdams, Julie A. Learn, Kermit L. Carraway

## Abstract

**Background:** Mucin-4 (MUC4) contributes to tumor cell survival and metastatic behavior, but its expression in breast cancer is unusual; previous studies have demonstrated that MUC4 levels are reduced in primary tumors relative to normal epithelium but are partially restored in metastatic lesions. Because DNA methylation is known to regulate *MUC4* gene expression in cultured cancer cells, we investigated whether methylation of the *MUC4* regulatory region changes across breast cancer progression.

**Results:** Analysis of the TCGA-BRCA database revealed that primary breast tumors exhibit increased methylation at CpG sites within the *MUC4* regulatory region relative to normal breast tissue, and methylation at key sites is inversely associated with *MUC4* transcript abundance. Moreover, DNA methyltransferase inhibition with 5-aza-2′-deoxycytidine alters *MUC4* expression in several breast cancer cell lines. Finally, targeted bisulfite sequencing analysis of an independent cohort of normal breast, primary breast tumor, and metastatic breast cancer specimens revealed that *MUC4* in primary tumors is significantly more methylated than in normal breast tissue at all 11 CpG sites examined. Metastatic specimens in this cohort generally display lower methylation than primary tumors and an intermediate methylation profile between normal and primary tissue.

**Conclusions:** Taken together, our findings support *MUC4* regulatory region methylation as a contributor to stage-dependent *MUC4* regulation.

## Background

Mucin-4 (MUC4) is a large membrane-associated glycoprotein that contributes to epithelial surface lubrication and protection but can also promote malignant behavior when aberrantly expressed. In cancer cells, MUC4 protein can alter cell-cell and cell-extracellular matrix interactions (1,2), promote resistance to cell death, facilitate immune evasion (3,4), and interact with ErbB2/HER2 to enhance pro-survival signaling and therapeutic resistance (5,6). Consistent with these functions, elevated *MUC4* expression has been associated with aggressive behavior in several epithelial malignancies (7). In breast cancer, however, *MUC4* displays an unusual pattern of expression. Previous studies from our laboratory have demonstrated that MUC4 protein is reduced in primary breast tumors relative to normal mammary epithelium but is partially restored in lymph node metastatic lesions (8). In an ErbB2 (HER2)-driven mouse mammary tumor model, MUC4 loss markedly reduced metastatic efficiency without altering primary tumor growth (9,10). Together, these observations suggest that *MUC4* may be regulated differently in primary and metastatic disease and raise the possibility that stage-dependent control of *MUC4* contributes to breast cancer progression.

Epigenetic remodeling is a common feature of breast tumorigenesis and metastasis, and DNA methylation can provide a stable but reversible mechanism for altering gene expression (11,12). *MUC4* is known to be subject to epigenetic regulation. Vincent et al. identified increased methylation near the *MUC4* transcriptional regulatory region in low-*MUC4*-expressing cancer cell lines and showed that reducing DNA methyltransferase activity could increase *MUC4* expression (13). Yamada et al. similarly mapped methylation across the 5′ region of *MUC4* and found that low-expressing cancer cell lines tended to exhibit higher methylation near the transcriptional regulatory region; treatment with the DNA methyltransferase inhibitor 5-aza-2′-deoxycytidine also increased *MUC4* expression in several cell lines (14). These studies have established a relationship between *MUC4* methylation and transcriptional output in cancer models, but whether *MUC4* methylation changes across stages of breast cancer progression has not been established.

We therefore hypothesized that methylation of the *MUC4* regulatory region changes during breast cancer progression in a manner consistent with the previously observed stage-specific pattern of MUC4 expression (8). To test this hypothesis, we investigated the methylation of the *MUC4* promoter region using retrospective patient data, breast cancer cell lines, and patient samples. We first assessed whether methylation within the *MUC4* regulatory region differs between primary breast tumors and normal breast tissue and whether methylation is associated with *MUC4* transcript abundance using the TCGA-BRCA database. We then examined the effect of pharmacologic DNA methyltransferase inhibition on *MUC4* expression in breast cancer cell lines. Finally, we used targeted bisulfite sequencing to evaluate methylation across the *MUC4* 5′ regulatory region in patient-derived normal, primary, and metastatic tissues. Together, these complementary approaches were employed to assess whether stage-associated changes in *MUC4* methylation are evident during breast cancer progression and are consistent with a role for DNA methylation in *MUC4* regulation.

## Materials and Methods

### Cell culture and reagents

JIMT-1, MDA-MB-231, MDA-MB-468, and MCF7 cell lines were purchased from the American Type Cell Collection (ATCC), authenticated by short-tandem repeat profiling (Genetics Core Facility at University of Arizona), and tested for mycoplasma contamination using PCR before use in the outlined studies (15). These cell lines were cultured in Dulbecco’s Modified Eagle Medium (DMEM) with L-Glutamine (Gibco) supplemented with 10% fetal bovine serum (Genesee) and 1% penicillin-streptomycin (Genesee) and were maintained at 37°C in 10% CO_2_.

### RT-QPCR

Cells were treated with 5µM 5-aza-2’-deoxycytidine (Sigma) or vehicle for 48 hours prior to RNA extraction for RT-QPCR. Total RNA was isolated using a PureLink RNA MiniKit (Ambion) and treated with the TURBO DNase kit (Invitrogen). Reverse transcription and qRT-PCR were performed using the Luna Universal One-Step RT-qPCR Kit (NEB) in a Bio-Rad iCycler CFX-96 real-time PCR machine. Transcript levels for human *MUC4* (Primers: 5’-CGCCATTGCTTTTGGGAGAC-3’, 5’-GCACCTGCGCCTATGGATAA-3’) were analyzed using Bio-Rad CFX Manager software and normalized to human glyceraldehyde-3-phosphate dehydrogenase expression (*GAPDH*; Primers: 5’-CGGAGTCAACGGATTTGGTC-3’, 5’-TTCTCAGCCTTGACGGTGC-3’) using the ΔΔCt method.

### TCGA data analysis

DNA methylation (Illumina Infinium HumanMethylation450 BeadChip, hg19) and RNA-seq expression data (log2[FPKM-UQ+1]) for *MUC4* were obtained from The Cancer Genome Atlas (TCGA) via MEXPRESS (16,17). Six CpG probes surrounding the *MUC4* transcriptional start site (TSS; cg13002712, cg08493265, cg10593292, cg19983801, cg07399928, cg01999399) were selected for analysis. Genomic coordinates for each probe were obtained from the Illumina 450K manifest annotation (IlluminaHumanMethylation450kanno.ilmn12.hg19, Bioconductor). Samples were classified as tumor or normal tissue based on the sample-type code embedded in the TCGA barcode (01 = primary tumor, n = 780; 11 = solid tissue normal, n = 84).

For each CpG site, methylation beta values were compared between tumor and normal samples using a two-sided Wilcoxon rank-sum test (Mann-Whitney U test), given the non-normal, bounded distribution typical of methylation beta values. Resulting *P*-values across the six CpG sites were adjusted for multiple comparisons using the Benjamini-Hochberg method to control the false discovery rate (FDR). Median beta values and their interquartile range were plotted for each CpG site.

The relationship between methylation and *MUC4* expression at each CpG site was assessed using Pearson correlation across all tumor and normal samples combined. For each CpG, methylation beta values and log2(FPKM-UQ+1)–transformed expression values were fit with an ordinary least squares linear regression model, and Pearson’s correlation coefficient (r) and associated two-sided p-value (testing the null hypothesis of no linear relationship) were calculated. Analyses were performed independently per CpG site. All analyses were performed in R (version 4.5.1) using the packages tidyr, dplyr, stringr, ggplot2, ggrepel, ggpubr, and minfi, along with the Bioconductor annotation package IlluminaHumanMethylation450kanno.ilmn12.hg19.

### Patient tissue processing and nucleic acid extraction

This study was reviewed by the UC Davis Institutional Review Board and was determined not to constitute human subjects research because it involved the use of de-identified, banked tissue specimens. All tissues were acquired from the UC Davis Biorepository Shared Resource after approval from the Biorepository Utilization Committee. Tissues were stored at −80°C prior to use and processed on dry ice up until lysis.

For simultaneous nucleic acid extraction, small pieces of frozen tissue (approximately 5mm x 5mm) were cut using a sterile chilled scalpel and added to cold lysis buffer containing β-mercaptoethanol (PureLink RNA Mini Kit, Invitrogen) before TissueRuptor homogenization (Qiagen, 2 x 30 second pulses full speed on ice). Lysates were passed 3x through a sterile 25g syringe before being centrifuged at 5000g for 5 minutes. The clear portion of the supernatant (avoiding the fatty layer) was transferred to a new tube and spun again at 12,000xg for 5 minutes before transferring the final supernatant to a DNA column (Qiamp Fast DNA Tissue Kit, Qiagen). The column was spun at top speed for 1min, and the flow-through was mixed at a 1:1 ratio with fresh 70% ethanol before transfer to an RNA column. The DNA column was stored on ice for later processing while the RNA was extracted according to normal kit instructions (PureLink RNA Mini Kit). DNA was then extracted according to normal kit instructions (Qiamp Fast DNA Tissue Kit). RNA samples were DNase treated using the TURBO DNase kit according to manufacturer instructions (Invitrogen). DNA samples were assessed for RNA contamination by Qubit (Invitrogen) and found to contain no RNA, so RNase digestion was not performed. All samples were stored at −80°C prior to downstream applications.

### RNA sequencing

RNA samples were sent to Plasmidsaurus (South San Francisco, CA) on dry ice for 3’ RNA-sequencing (RNA-seq). There, sequencing libraries were generated by converting mRNA into complementary DNA (cDNA) via reverse transcription and second-strand synthesis, followed by tagmentation, library indexing, and amplification. Libraries were sequenced on the Illumina platform using a 3’ end counting approach, targeting approximately 10 million deduplicated reads (20 million raw reads) per sample.

The quality of raw FASTQ files was assessed using FastQC (v0.12.1). Reads were quality-filtered using fastp (v0.24.0), applying poly-X tail trimming, 3’ quality-based tail trimming, a minimum Phred quality score of 15, and a minimum read length of 50 bp. Quality-filtered reads were aligned to the human reference genome (GRCh38) using STAR (v2.7.11) with non-canonical splice junction removal, and unmapped reads were retained in output. Aligned reads were coordinate-sorted using Samtools (v1.22.1) and PCR and optical duplicates were removed via UMI-based deduplication using UMIcollapse (v1.1.0). Alignment quality metrics, strand specificity, and read distribution across genomic features were assessed using RSeQC (v5.0.4) and Qualimap (v2.3). Gene-level expression was quantified using featureCounts (Subread package, v2.1.1) with strand-specific counting and fractional assignment of multi-mapping reads. Exons and 3’ UTRs were used as feature identifiers, grouped by gene ID.

To investigate low gene counts at the *MUC4* locus, reads overlapping *MUC4* (GRCh38, chr3:195,746,765–195,811,973, minus strand) were extracted from BAM files using Rsamtools/GenomicAlignments in R and inspected for MAPQ value. Additionally, reads flagged as unmapped by STAR were extracted using Samtools bam2fq and realigned against the *MUC4* genomic sequence (Ensembl, GRCh38) using minimap2 in splice-aware mode, which permits gapped alignment across intron-spanning reads not achievable with the sequence alone. Rescued reads (i.e., those aligning to the *MUC4* locus despite being classified as unmapped by STAR) were characterized by mapping quality (MAPQ), CIGAR string (to identify spliced alignments), and genomic position relative to the gene’s annotated 3’ end.

For sample clustering (Supplementary Figure S1), normalized expression values (counts per million, CPM) were filtered to remove low-expression genes, retaining only genes with CPM > 1 in at least 3 samples. Filtered CPM values were log2-transformed (log2[CPM + 1]) and samples were hierarchically clustered using “ward.D2” linkage; results were visualized as a heatmap with Pheatmap (v1.0.13) in R. Gene set enrichment analysis (GSEA) was performed for each of the three pairwise comparisons against the Hallmark Human 2025 gene set collection from MSigDB using GSEApy (v0.12). Enrichment significance was assessed using a false discovery rate (FDR)-corrected q-value, with normalized enrichment score (NES) reported as the primary effect-size measure. For visualization, the top 20 gene sets by minimum FDR q-value across the three comparisons were selected and displayed in a dot plot, with dot color representing NES and dot size representing statistical significance (−log10 FDR q-value).

### Tumor purity analysis

For primary tumor and metastasis samples, sample tumor fraction was estimated using PUREE (v1.0), a linear regression model trained via weakly supervised learning on rank-percentile-transformed gene expression profiles and matched genomic consensus purity estimates from 7,864 TCGA solid tumor samples spanning 20 cancer types (18). PUREE was run in Python using pinned dependency versions (scikit-learn 1.1.2, numpy 1.23.2, pandas 1.4.3, joblib 1.1.0). As PUREE was developed and validated on whole-transcript RNA-seq rather than the 3’ end counting protocol used to generate this dataset, and no independent (e.g., histological) purity assessment was available for these samples, absolute purity values should be interpreted with caution; estimates are presented primarily to support relative comparison of tumor cellularity across samples within this study. Tumor samples with an estimated tumor fraction of 0.4 or less were excluded from further analysis, consistent with purity thresholds used elsewhere (19,20).

### Targeted bisulfite sequencing

Genomic DNA was sent to CD Genomics (Shirley, NY) on dry ice for targeted bisulfite sequencing. There, 1ug of DNA was converted using the EZ DNA Methylation-Gold Kit (Zymo) before PCR amplification with primers designed to cover a 1kb region surrounding the *MUC4* TSS (see Supplementary Table 1) using the MethPrimer software (21).

PCR products were pooled and barcoded before sequencing on the Illumina Hiseq platform. Sequencing data were quality filtered using the Trimmomatic-0.36 software and alignment was performed using Bsmap (v2.73) with default parameters. Methylated cytosines were called using a binomial distribution as described by Lister et al. (22) and methylation levels were defined as the fraction of read counts of ‘C’ in the total read counts of both ‘C’ and ‘T’ for each covered C site. Only those CpGs covered by at least 200 reads in at least one sample were considered for testing.

For each of the 11 CpG sites detected during sequencing, an omnibus Kruskal-Wallis test was used to assess whether percent methylation differed across the three tissue types (Normal, Primary, Metastasis); for sites with a significant omnibus result (FDR < 0.05), pairwise comparisons between tissue types were performed using Dunn’s test with Benjamini-Hochberg correction.

## Results

### MUC4 regulatory region methylation differs between normal breast tissue and primary breast tumors in TCGA

To determine whether *MUC4* methylation differs between normal breast tissue and primary breast tumors, we analyzed DNA methylation and RNA-seq data from the TCGA-BRCA cohort obtained through MEXPRESS (16,17). Six CpG probes within the *MUC4* 5′ regulatory region were examined in primary breast tumors and normal breast tissue. At the two most proximal CpG sites, primary tumors were significantly more highly methylated than normal breast tissue.

We next assessed whether methylation at these sites was associated with *MUC4* transcript abundance. Methylation was inversely correlated with *MUC4* expression, although the magnitude of the relationship was modest. These findings identify an association between increased *MUC4* regulatory-region methylation and reduced *MUC4* transcript abundance in primary breast tumors (Fig. 1).

**Figure 1.**
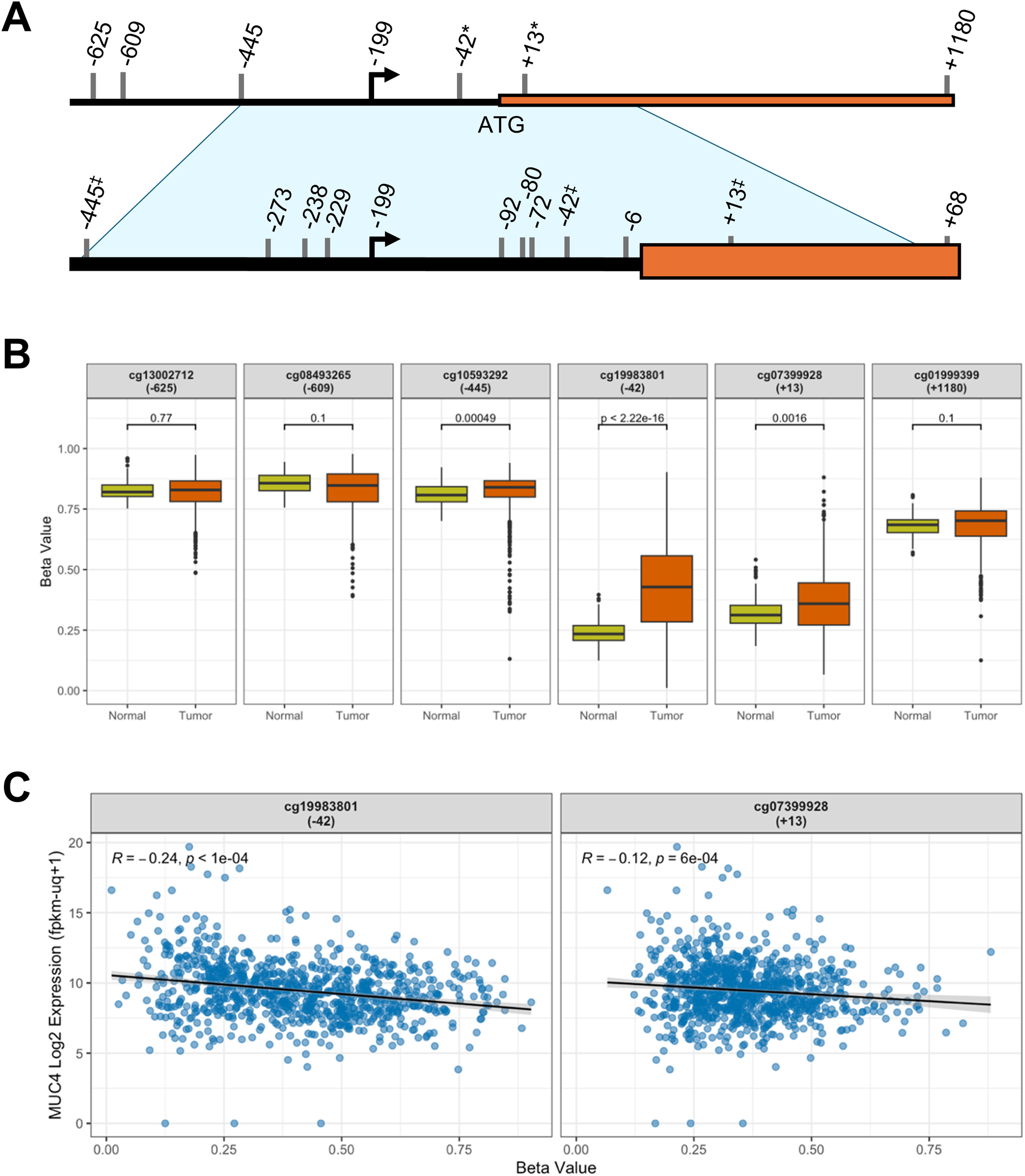
*MUC4* promoter methylation is increased in primary breast tumors and inversely associated with expression. *MUC4* expression data (log2[FPKM-UQ+1]) and DNA methylation beta values for breast invasive carcinoma and adjacent normal samples were obtained from The Cancer Genome Atlas (TCGA-BRCA) via MEXPRESS (mexpress.be; ref 17). **A)** CpG sites surrounding the *MUC4* TSS (−199 relative to start codon) are depicted. *Top*, sites captured in the TCGA methylation microarray (panel B); *bottom*, sites assayed by targeted bisulfite sequencing (Figure 3). *, sites tested for methylation vs transcript expression correlation (panel C); ^‡^, sites captured in both microarray and targeted bisulfite sequencing. **B)** Comparison of methylation beta value at six sites flanking the *MUC4* start codon for normal mammary tissue (n = 84) vs breast invasive carcinoma (n = 780) samples. Distance from the start codon is indicated in each plot title. *P*-values generated by Wilcoxon rank-sum test with Benjamini Hochberg correction; box plots depict median and quartiles. **C)** Pearson correlation between *MUC4* transcript expression and methylation in TCGA samples from panel B. Solid lines represent linear regression fits; shaded areas indicate 95% confidence intervals.

### DNA methyltransferase inhibition alters MUC4 expression in breast cancer cell lines

Previous studies demonstrated that *MUC4* expression can be influenced by DNA methylation in cancer cells. To determine whether *MUC4* expression is similarly responsive to DNA methyltransferase inhibition in breast cancer cells, MCF7, JIMT-1, MDA-MB-468, and MDA-MB-231 cells were treated with 5-aza-2′-deoxycytidine for 48h and MUC4 transcript levels were measured by RT-qPCR. *MUC4* expression increased following 5-Aza treatment in three of the four cell lines examined. The largest response was observed in JIMT-1 cells, in which *MUC4* transcript levels increased approximately 15-fold. MDA-MB-468 cells showed an approximately 50% increase in *MUC4* expression, whereas MDA-MB-231 cells showed an approximately 50% decrease. Thus, *MUC4* expression was responsive to DNMT inhibition in multiple breast cancer cell lines, although the direction and magnitude of the response were cell-line dependent (Fig. 2).

**Figure 2.**
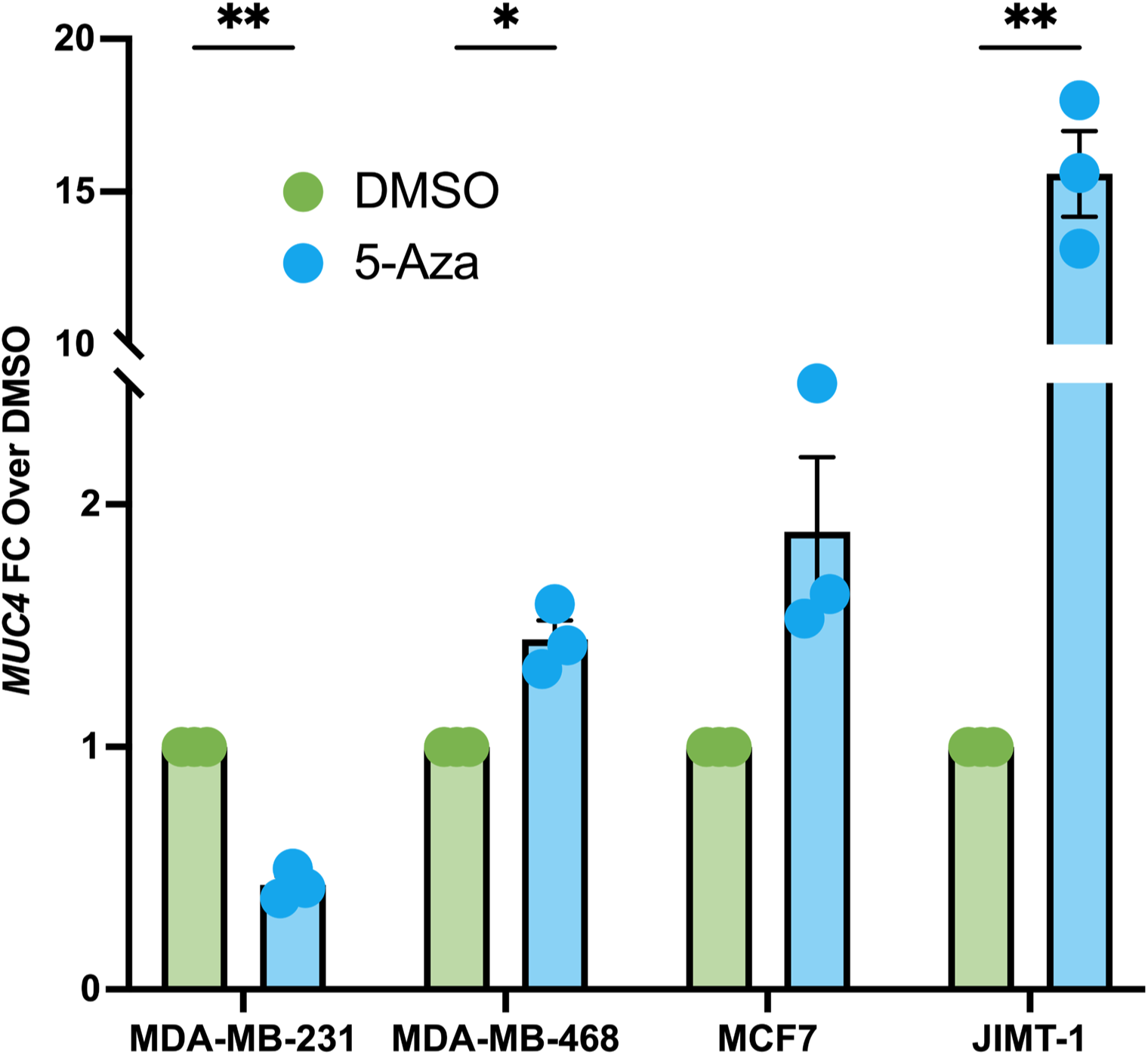
DNA methyltransferase inhibition alters MUC4 expression in breast cancer cells. Breast cancer cell lines were treated with DMSO vehicle or 5µM 5-aza-2’-deoxycytidine (5-Aza) for 48hrs prior to RT-QPCR. Transcript abundance was normalized to *GAPDH* using the ΔΔCt method. Error bars represent SEM, n=3. Statistical significance was calculated using an unpaired two-tailed t-test with Welch’s correction (*, *P* < 0.05, **, *P* < 0.01).

### MUC4 methylation differs among normal breast, primary tumor, and metastatic breast cancer tissues

Because TCGA-BRCA primarily enables comparison of normal breast tissue with primary breast tumors, we next examined *MUC4* methylation across normal, primary, and metastatic breast tissues using an independent UC Davis biorepository cohort. Twelve specimens from each tissue group were initially obtained (see Table 1). DNA and RNA were isolated from the same tissue specimens, and targeted bisulfite sequencing was performed across a 1-kb region surrounding the *MUC4* 5′ regulatory region. This region was chosen because it covers the *MUC4* proximal promoter (−219/-1) and transcription initiation site (−199), as well as the region defined by Yamada et al. (14) to be most closely correlated to *MUC4* gene expression.

**Table 1.** Patient information for *MUC4* methylation analysis. Characteristics of patient tissue samples, accessed via the UC Davis Biorepository Shared Resource.

| **Tissue Type (n)** | **Average Age (y)** | **Location** | **Diagnosis** |
|---|---|---|---|
| Non-Malignant (12) | 54.9 | Breast | N/A |
| Primary Tumor (12) | 52.1 | Breast | Ductal carcinoma (11) |
|  |  |  | Ductal carcinoma in situ (1) |
| Metastasis (12) | 48.5 | Axillary lymph node (4) | Breast carcinoma |
|  |  | Femur (2) |  |
|  |  | Brain (5) |  |
|  |  | Ovary (1) |  |
| Total (36) | 51.8 |  |  |

Because histologic tumor-purity estimates were unavailable for the frozen specimens, tumor cellularity was estimated from 3′ RNA-seq data using PUREE. Samples with an estimated tumor fraction of 0.4 or less, or samples without adequate RNA-seq data, were excluded from methylation analysis. Following filtering, the methylation cohort included 11 normal breast samples, 5 primary tumors, and 10 metastatic specimens.

Targeted bisulfite sequencing yielded 11 CpG sites meeting the prespecified coverage criteria, three of which overlapped CpGs represented on the TCGA 450K array. Primary breast tumors were significantly more methylated than normal breast tissue at all 11 CpG sites examined. Metastatic samples were also significantly more methylated than normal tissue at nine of the 11 sites. Across most CpGs, however, methylation in metastatic specimens was lower than in primary tumors, resulting in an overall intermediate methylation pattern. Following correction for multiple comparisons, two CpG sites remained significantly different between primary and metastatic tumors (Fig. 3B).

**Figure 3.**
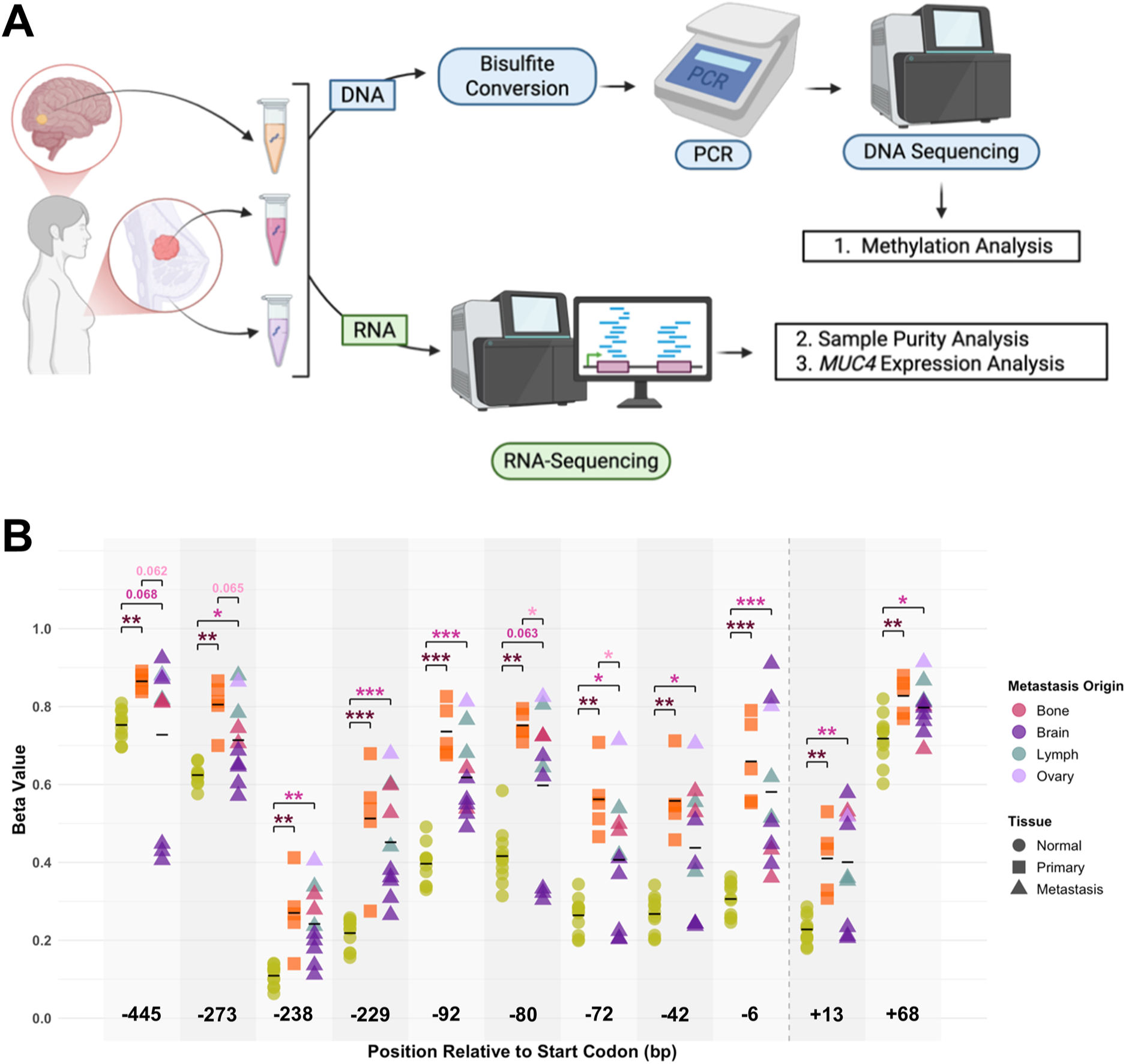
*MUC4* is differentially methylated across breast cancer progression in patient samples. **A)** Schematic of experimental approach. DNA and RNA were simultaneously extracted from patient samples. DNA was bisulfite treated, *MUC4* was PCR amplified and sequenced, and methylation status was assessed (CD Genomics, Shirley, NY). RNA was 3’ sequenced for differential *MUC4* expression analysis (Plasmidsaurus, South San Francisco, CA) and sample tumor purity analysis. Figure created with BioRender. **B)** Percent methylation was measured by targeted bisulfite sequencing at 11 CpG sites positioned relative to the start codon (dashed line). Each point represents a sample, black lines represent tissue mean (Normal, n = 11; Primary, n = 5; Metastasis, n = 10), and metastatic tissue of origin indicated by color. Statistical comparisons across tissue types (Kruskal-Wallis with Dunn’s post-hoc test, BH-FDR corrected) are reported above each CpG site (*, p.adj < 0.05; **, p.adj < 0.01; ***, p.adj < 0.001).

Exploratory comparison of metastatic specimens by tissue site suggested heterogeneity in methylation patterns, with brain metastases tending toward lower methylation across several CpG sites. However, the limited number of samples within individual metastatic sites precluded formal site-specific statistical analysis (Fig. 3C).

We also sought to assess the relationship between *MUC4* methylation and transcript abundance within the independent tissue cohort. However, *MUC4* gene-level counts from 3′ RNA-seq were insufficient for reliable differential expression analysis. Locus-level inspection and realignment of reads initially classified as unmapped recovered a small number of additional *MUC4*-compatible reads per sample, but overall coverage remained insufficient for quantitative comparison among tissue groups. *MUC4* expression was therefore not analyzed further in this cohort.

## Discussion

MUC4 is an established contributor to metastatic behavior, with reported anti-adhesive (1), anti-anoikis (9), and pro-migratory properties (23,24), and it promotes efficient metastasis *in vivo* (9). In breast cancer, however, MUC4 expression appears to be dynamically regulated across disease progression. Previous work from our laboratory showed that MUC4 protein expression is reduced in primary breast tumors relative to normal adjacent tissue but partially restored in matched lymph node metastases (8). In a HER2-driven mammary tumor model, loss of MUC4 markedly reduced metastatic efficiency without altering primary tumor growth (9). These observations raise the possibility that the biological requirements for MUC4 differ between primary and metastatic disease. In the present study, we identify stage-associated changes in *MUC4* regulatory-region methylation that are consistent with this previously described expression pattern. Primary tumors showed increased *MUC4* methylation relative to normal breast tissue, whereas metastatic specimens generally displayed an intermediate methylation profile. Together with the inverse relationship between methylation and MUC4 transcript abundance in TCGA and the responsiveness of *MUC4* expression to DNA methyltransferase inhibition, these findings support DNA methylation as one component of stage-dependent *MUC4* regulation.

*MUC4* is a large and transcriptionally complex gene whose expression is regulated through multiple mechanisms (7). Previous studies specifically implicated DNA methylation in this regulation. Vincent et al. demonstrated that epigenetic control of *MUC4* involves both DNA methylation and histone modification and showed that perturbation of DNA methyltransferase activity can alter *MUC4* expression (13). Yamada et al. similarly identified an association between CpG methylation in the *MUC4* 5′ regulatory region and *MUC4* expression across cancer cell lines and showed that treatment with 5-aza-2′-deoxycytidine could increase *MUC4* expression (14). We extended these observations to a panel of breast cancer cell lines. *MUC4* transcript abundance increased following 5-Aza treatment in three of four lines, with the largest response observed in JIMT-1 cells. However, MDA-MB-231 cells showed reduced *MUC4* expression following treatment. This heterogeneity suggests that DNA methylation is unlikely to act as the sole determinant of *MUC4* expression and that its influence is dependent on cellular context. Moreover, because 5-Aza alters DNA methylation genome-wide, these experiments do not establish that the observed transcriptional changes result from direct demethylation of the *MUC4* locus.

Analysis of TCGA-BRCA provided independent evidence linking *MUC4* methylation and transcription in human breast tissue. Primary breast tumors showed increased methylation relative to normal breast tissue at CpG sites within the proximal *MUC4* regulatory region, and methylation at key sites was inversely associated with *MUC4* transcript abundance. Although the magnitude of these correlations was modest, such a relationship is consistent with DNA methylation contributing to *MUC4* regulation as one component of a broader transcriptional and epigenetic program. The relatively broad distribution of methylation values among primary tumors also suggests considerable biological heterogeneity. Such heterogeneity could reflect differences in breast cancer subtype, transcriptional state, tumor microenvironment, or other characteristics, although the present study was not designed or powered to identify the basis of this variation.

The independent patient cohort extended these observations across normal breast tissue, primary breast tumors, and metastatic breast cancer. Primary tumors were significantly more methylated than normal breast tissue at all 11 CpG sites examined, providing evidence that increased methylation across this MUC4 regulatory region accompanies primary tumorigenesis. Metastatic specimens remained more methylated than normal breast tissue at most sites but generally showed lower methylation than primary tumors, producing an intermediate methylation profile. Although only two CpG sites remained significantly different between primary and metastatic tumors following correction for multiple comparisons, the direction of the change was broadly consistent across the assayed region. These findings are compatible with partial loss of primary-tumor-associated *MUC4* methylation during metastatic progression.

The metastatic specimens represented both lymphatic and distant metastatic sites, including lymph node, bone, brain, and ovary. Exploratory visualization suggested some heterogeneity among metastatic sites, with brain metastases tending toward lower *MUC4* methylation across several CpGs. However, the number of specimens from each site was too small to permit meaningful site-specific statistical analysis. Accordingly, these observations should be considered hypothesis-generating rather than evidence for organ-specific *MUC4* methylation patterns. Larger cohorts that permit stratification by metastatic site and breast cancer subtype will be required to determine whether the degree of *MUC4* methylation remodeling differs among biologically distinct metastatic contexts.

Importantly, the present findings establish association rather than causation. The study does not directly demonstrate that loss of methylation at specific *MUC4* CpGs causes re-expression of MUC4 during metastatic progression, nor does it show that methylation changes at the *MUC4* locus directly alter metastatic behavior. Rather, three complementary observations support a contribution of DNA methylation to *MUC4* regulation: *MUC4* regulatory region methylation is inversely associated with transcript abundance in TCGA; *MUC4* expression is responsive to DNMT inhibition in breast cancer cells; and *MUC4* methylation changes across normal, primary, and metastatic patient tissues. Taken together, these findings support a model in which DNA methylation participates in the dynamic regulation of *MUC4* during breast cancer progression but functions alongside other transcriptional and epigenetic mechanisms.

Several limitations should be considered. First, the independent tissue cohort was modest in size, particularly after tumor-purity filtering, which limited statistical power for primary-versus-metastatic comparisons and precluded robust analysis by molecular subtype or metastatic site. Second, tumor purity was estimated computationally from 3′ RNA-seq rather than by histopathologic assessment. PUREE was developed and validated using whole-transcript RNA-seq rather than the 3′ end-counting strategy used here (18), and absolute purity estimates should therefore be interpreted cautiously. Third, the 5-Aza experiments provide indirect evidence for methylation-dependent regulation because the treatment affects DNA methylation throughout the genome rather than selectively at *MUC4*. Finally, *MUC4* transcript abundance could not be reliably quantified in the same patient specimens used for targeted bisulfite sequencing, preventing direct within-sample analysis of methylation and expression. Although locus-level inspection and realignment recovered a small number of additional *MUC4*-compatible reads, coverage remained insufficient for quantitative comparison. The complexity of the *MUC4* transcript architecture, including extensive alternative splicing and 3′ transcript diversity (25–27), may contribute to the difficulty of quantifying *MUC4* using a 3′ end-counting approach, although the present data cannot establish the reason for the low read counts.

Future studies could directly test the functional importance of the methylation changes identified here. Expanded tissue cohorts would permit assessment of breast cancer subtype and metastatic-site-specific patterns, while paired methylation and robust transcript or protein measurements could establish whether methylation changes correlate with *MUC4* expression within individual specimens. In addition, locus-specific epigenome-editing approaches now make it possible to manipulate methylation at defined genomic regions without the genome-wide effects of pharmacologic DNMT inhibition (28). Such approaches could determine whether individual CpGs or regulatory regions within *MUC4* directly control transcription and whether these changes influence phenotypes relevant to metastatic progression.

## Conclusions

This study identifies stage-associated changes in *MUC4* regulatory-region methylation during breast cancer progression. Primary tumors show increased methylation relative to normal breast tissue, whereas metastatic specimens generally exhibit partial loss of this primary-tumor-associated methylation pattern. Together with the inverse methylation-expression relationship observed in TCGA and the responsiveness of *MUC4* expression to DNMT inhibition, these findings support DNA methylation as one contributor to stage-dependent *MUC4* regulation. Direct locus-specific studies will be required to determine whether these methylation changes are causal.

## Declarations

### Ethics approval and consent to participate

This study was reviewed by the UC Davis Institutional Review Board and was determined not to constitute human subjects research because it involved the use of de-identified, banked tissue specimens. All tissues were acquired from the UC Davis Biorepository Shared Resource after approval from the Biorepository Utilization Committee.

### Consent for publication

Not applicable

### Availability of data and materials

The TCGA dataset supporting the conclusions of this article is publicly available via the NCI Genomic Data Commons (https://portal.gdc.cancer.gov/), project TCGA-BRCA. Data were accessed and visualized using MEXPRESS (https://mexpress.be; accessed June 2026).

### Authors’ Contributions

SF and KLC contributed to conceptualization; SF and KLC contributed to methodology; SF, LLS, and MM contributed to data collection; SF and JL contributed to data analysis and interpretation; KLC contributed resources; SF and KLC contributed to manuscript writing and review. All authors read and approved the final manuscript.

## Acknowledgements

Thank you to the UC Davis Comprehensive Cancer Center Genomics Shared Resource (for data analysis support) and Biorepository Shared Resource (for tissue acquisition support), both supported by NIH grant P30CA93373.

