## Supplementary Figures for "*MUC4* regulatory-region methylation is remodeled during breast cancer progression and metastasis"

| Primers | |
|---|---|
| Set 1 | F: 5’-ATATGGTTTTTYGTTTTTTAATTTTTTGGGGTTTG-3’<br>R: 5’-TATCCCACCAAAACTTTAAACTACTAAA-3’ |
| Set 2 | F: 5’-GGAGAGGATTTAGGAATTYGGTTAGGA-3’<br>R: 5’-TTCTCTCCTCAACCCCACCCCACTAC-3’ |
| Set 3 | F: 5’-GGAGAAAYGTATTTGGTTYGGGTTAGT-3’<br>R: 5’-TCCATCACTTACCTAAAACCACATAC-3’ |

**Supplementary Table 1. *MUC4* PCR primers.**

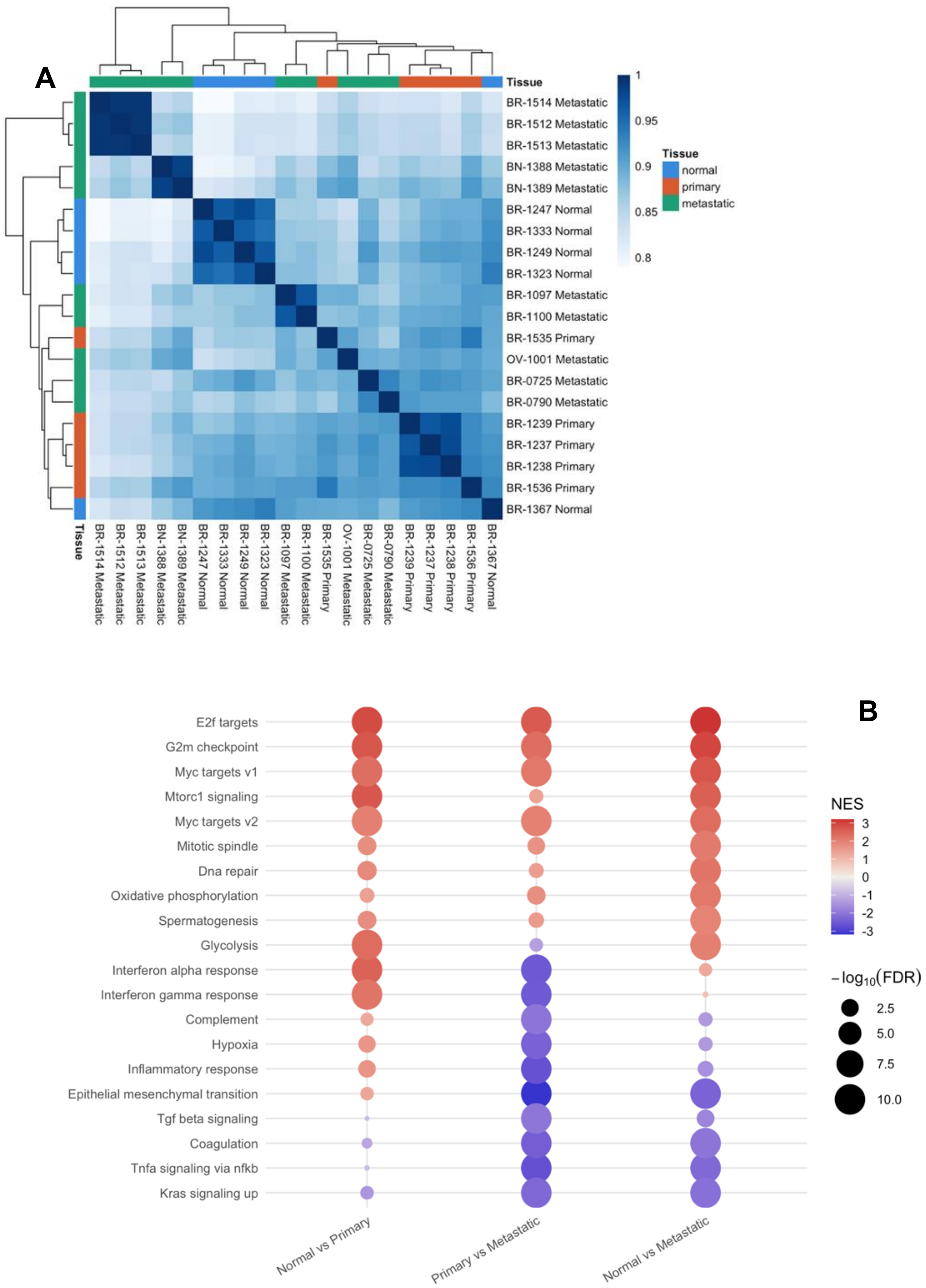
A
Tissue
BR-1514 Metastatic
BR-1512 Metastatic
BR-1513 Metastatic
BN-1388 Metastatic
BN-1389 Metastatic
BR-1247 Normal
BR-1333 Normal
BR-1249 Normal
BR-1323 Normal
BR-1097 Metastatic
BR-1100 Metastatic
BR-1535 Primary
OV-1001 Metastatic
BR-0725 Metastatic
BR-0790 Metastatic
BR-1239 Primary
BR-1237 Primary
BR-1238 Primary
BR-1536 Primary
BR-1367 Normal
1
0.95
0.9
0.85
0.8
Tissue
normal
primary
metastatic
B
E2f targets
G2m checkpoint
Myc targets v1
Mtorc1 signaling
Myc targets v2
Mitotic spindle
Dna repair
Oxidative phosphorylation
Spermatogenesis
Glycolysis
Interferon alpha response
Interferon gamma response
Complement
Hypoxia
Inflammatory response
Epithelial mesenchymal transition
Tgf beta signaling
Coagulation
Tnfa signaling via nfkb
Kras signaling up
Normal vs Primary
Primary vs Metastatic
Normal vs Metastatic
NES
3
2
1
0
-1
-2
-3
$-\log_{10}(FDR)$
2.5
5.0
7.5
10.0

**Supplementary Figure 1. Sample clustering and gene program differences between normal breast, primary tumor, and metastatic tissue. A)** Hierarchical clustering of RNAseq samples by pairwise Pearson correlation of Log2 counts-per-million (CPM) values. Color scale reflects the Pearson correlation coefficient between each sample pair, from low (white) to high (blue). Tissue type (normal, primary, metastatic) is indicated by the annotation bar along the top and side of the heatmap. **B)** Gene set enrichment analysis (GSEA) was performed in the Plasmidsaurus interface for each pairwise comparison (normal vs. primary, primary vs. metastatic, normal vs. metastatic) against the Hallmark Human 2025 gene set collection. Dot plot shows the top 20 enriched gene sets ranked by significance across the three comparisons. Dot color represents the normalized enrichment score (NES), with red indicating enrichment toward the second term (i.e., “Primary” in “Normal vs Primary”) and blue indicating enrichment toward the first term. Dot size corresponds to statistical significance (−log10 FDR q-value). Gene sets are ordered by mean NES across comparisons.

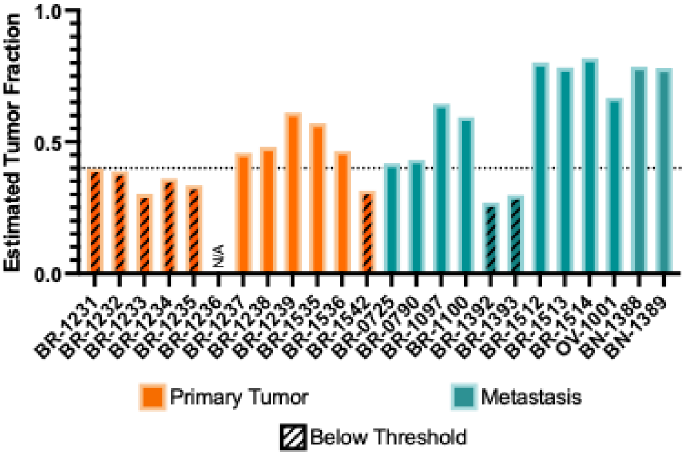


**Supplementary Figure 2. Tumor purity analysis using PUREE.** Tumor purity was estimated for each sample using PUREE, a pan-cancer linear regression model trained on whole-transcript RNA-seq data from The Cancer Genome Atlas (TCGA). (18). Samples below 0.4 (striped boxes) or samples that did not yield RNA sequencing results (N/A) were excluded from further analysis (19,20).

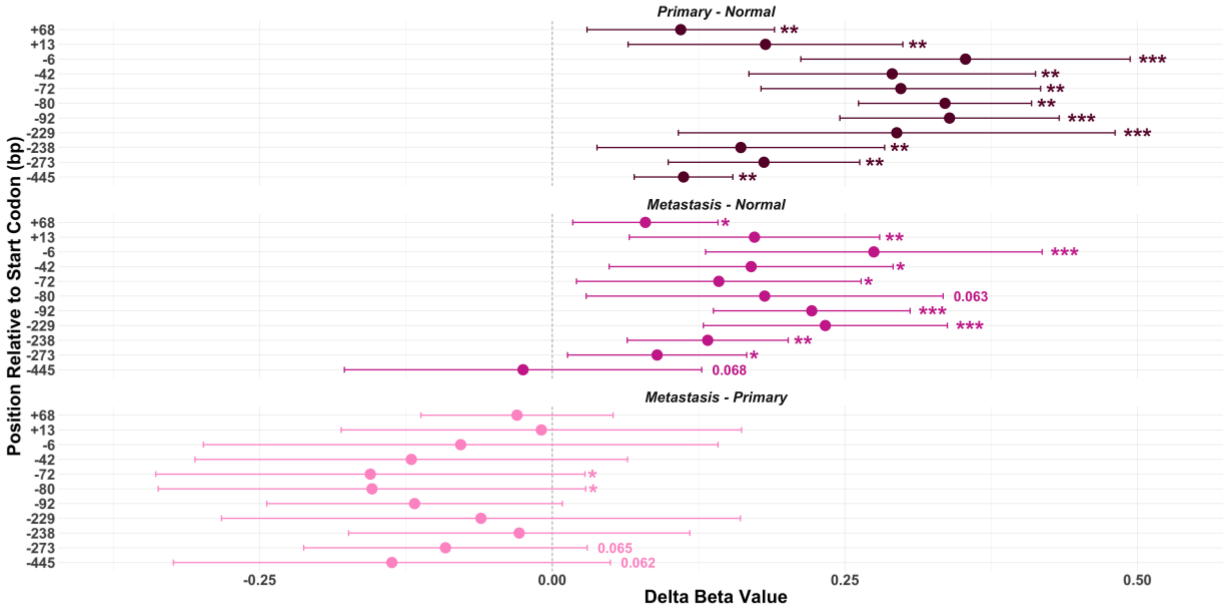


**Supplementary Figure 3. Patient differential methylation analysis.** For each CpG site, the difference in mean percent methylation (delta beta value) between two tissue types is shown as a point, with horizontal bars representing the 95% confidence interval of the difference. Statistical comparisons across tissue types (Kruskal-Wallis with Dunn's post-hoc test, BH-FDR corrected) are reported to the right each CpG site (*** = p.adj < 0.001, ** = p.adj < 0.01, * = p.adj < 0.05).
